# The IPF fibroblastic focus is an active collagen biosynthesis factory embedded in a distinct extracellular matrix

**DOI:** 10.1101/2021.11.06.467549

**Authors:** Jeremy A. Herrera, Lewis Dingle, M. Angeles Montero, Rajesh Shah, Rajamiyer V Venkateswaran, John F. Blaikley, Craig Lawless, Martin A. Schwartz

## Abstract

**Background:** The Fibroblastic Focus (FF) is the signature lesion of Idiopathic Pulmonary Fibrosis (IPF) where myofibroblasts accumulate and extracellular matrix (ECM) is produced. However, the molecular composition and function of the FF and surrounding tissue remain undefined.

**Methods:** Utilizing laser capture microdissection coupled mass spectrometry (LCM-MS), we interrogated the FF, adjacent mature scar, and adjacent alveoli in 6 IPF specimens plus 6 non-fibrotic alveolar specimens as controls. The data were subject to qualitative and quantitative analysis, and validation by immunohistochemistry.

**Results:** We found that the protein signature of IPF alveoli is defined by immune deregulation as the strongest category. The IPF mature scar was classified as end-stage fibrosis whereas the FF contained an overabundance of a distinctive ECM compared to non-fibrotic control.

**Conclusion:** Spatial proteomics demonstrated distinct protein compositions in the histologically defined regions of IPF tissue. These data revealed that the FF is the main site of collagen biosynthesis and that the alveoli adjacent to the FF are abnormal. This new and essential information will inform future mechanistic studies on mechanisms of IPF progression.

## Introduction

Idiopathic Pulmonary Fibrosis (IPF) is a progressive fibrotic lung disease characterized by the excessive deposition of extracellular matrix (ECM). IPF histology corresponds to the Usual Interstitial Pneumonia (UIP) pattern. UIP is a fibrotic lung disease, marked by sub-pleural and para-septal fibrosis alternating with patchy areas of morphologically normal lung and honeycomb pattern elsewhere [1]. The areas between fibrosis and morphologically normal lung typically contain fibroblastic growths of variable size termed the fibroblastic focus (FF). These structures are sites of myofibroblast accumulation within a pathological ECM, consistent with the notion that fibrosis spreads from the FF into uninvolved alveoli [2]. 3-dimensional reconstructions of the FF reveal both complex continuous structures and discrete lesions of variable shape [3, 4]. IPF is generally considered a fibro-proliferative disease, however, several reports show that the myofibroblasts within the FF are not proliferative [5–9]; suggesting that these myofibroblasts serve other biological functions.

It is proposed that FF form due to repetitive epithelial injury and damage responses leading to myofibroblast accumulation [10]. The origin of the myofibroblasts within the FF remains an active area of investigation. Available literature suggests that resident lung fibroblasts alone are insufficient, but that damaged and stressed alveolar epithelial cells transitioning into myofibroblasts via epithelial-to-mesenchymal transition (EMT) also contribute [11]. Progenitor cells (resident or circulating) may also differentiate into progeny myofibroblasts to promote these lesions [12–14]. A general consensus is that the FF and surrounding tissue is an invasive front where myofibroblasts contract, synthesize collagen, and progress towards adjacent uninvolved alveoli [2, 8]. Thus, a deeper understanding of the fibroblastic focus is needed to determine its biological functions and role in IPF progression.

Recently, our group has developed a laser capture microdissection-coupled mass spectrometry (LCM-MS) technique to characterize the ECM of fixed and stained human lung tissue [15], which we now apply to the FF and surrounding tissue. Here we present a comprehensive tissue atlas that defines cell surface and ECM proteins, and pathway analysis of histologically defined regions of the IPF lung in an effort to understand the pathogenesis of IPF.

## Materials and Methods

### Human lung tissue procurement

Patient-consented tissue was collected following University of Manchester Health Research Authority protocols: (REC#14/NW/0260) for IPF specimens and (REC#20/NW/0302) for non-fibrotic control specimens. The specimens were diagnosed as Idiopathic Pulmonary Fibrosis (IPF) as defined by current guidelines [1]. Non-fibrotic controls were collected from morphologically normal lung tissue distal to tumors during resection. In this study, we used 6 IPF and 6 non-fibrotic lung specimens.

### Histological staining

Human lung specimens were formalin-fixed and paraffin-embedded (FFPE) and sectioned at 5 μm on appropriate slides. H&E-staining was achieved using an automated stainer (Leica XL) at University of Manchester’s Histology Core. Briefly, the slides were heated at 60°C for 20 minutes, dewaxed with xylene, dehydrated by alcohol treatment, and rinsed in tap water. Slides were then hematoxylin-stained for 2 minutes, acid-alcohol treated, and stained for 1 minute in eosin. Slides were then washed in 100% ethanol and allowed to air-dry. These slides were stored at 4°C for up to one week while performing laser capture microdissection (LCM). For pentachrome staining, we used a modified Russel-Movats pentachrome stain protocol which we have previously described in detail [15].

For immunohistochemistry (IHC), we utilized the Novolink Polymer Detection Systems (Leica, RE7200-CE) as previously described in detail [16]. Deparaffinized and rehydrated 5-μm FFPE sections were subjected to antigen heat retrieval using citrate buffer (abcam, ab208572), in a pre-heated steam bath (100°C) for 20 minutes, before cooling to room temperature for 20 minutes. Slides were then treated with 3-4% hydrogen peroxide (Leica Biosystems RE7101) for 10 minutes, blocked in SuperBlock™ (TBS, Thermo Scientific™; 37581) buffer for a minimum of 1 hour, and probed with primary antibodies overnight at 4°C in 10% SuperBlock™ solution in Tris Buffered Saline Tween 20 solution (TBS-T, pH 7.6). Primary antibodies used were as follows: SerpinH1/HSP47 (abcam ab109117, titre 1:20,000), Collagen 12A1 (abcam ab21304, titre 1:200).

The following day, the specimens were subjected to Novolink Polymer Detection Systems (Leica Biosystems RE7270-RE, per the manufacturer’s recommendations), with multiple TBS-T washes. Sections were developed for 5 minutes with DAB Chromagen (Cell Signaling Technology^®^, 11724/5) before being counterstained with hematoxylin, dehydrated through sequential ethanol and xylene and cover-slipped with Permount mounting medium (Thermo Scientific™, SP15).

### Laser Capture Microdissection

To precisely perform this experiment, 3 serial sections were created per specimen. The first section was placed onto a standard glass slide for pentachrome stain, and the next two serial sections were placed onto two separate MMI membrane slides (MMI, 50102) for H&E staining (see above for staining details). The MMI CellCut Laser Microdissection System (Molecular Machines & Industries) can load up to 3 slides. In the first slot, we loaded the first pentachrome stained slide followed by the two H&E-stained MMI slides. We used MMI CellCut software to perform a complete CellScan of the pentachrome slide and H&E slides to allow for careful tissue registration. Using the pentachrome slide as a guide, we then used a closed-shape drawing tool to outline regions of interest. Automated cutting was achieved by adjusting the settings to a laser focus of 350 μm, a 60% laser power, and 50 μm/second speed. Using adhesive MMI transparent caps (MMI, 50204) and MMIs CapLift technology, we gently lift dissected specimens onto the adhesive caps which were stored in −20°C for several weeks until all samples were collected and processed for mass spectrometry. Ultimately, we used up to 30 sections (in sets of 3) to capture desired volumes (~0.1 mm^3^) per region of the IPF specimen.

### Mass spectrometry sample preparation

Samples were prepared using a multi-step method as previously described in our detailed technical report with minor modifications [15]. Firstly, after samples are trypsin digested, instead of eluting in 50% acetonitrile with 0.2 % formic acid, we now elute in 30% acetonitrile with 0.2% formic acid. In addition, after those samples are dried by speedvac, we now resuspend in 3% acetonitrile with 0.1% formic acid (instead of 5% acetonitrile with 0.1% formic acid). Lastly, our final elutions after de-salting our samples now uses 30% acetonitrile (instead of 50% acetonitrile). These were important as we began experiencing downstream contamination that we speculated was due to high acetonitrile concentrations; a problem that was resolved by lowering acetonitrile.

### Liquid chromatography coupled tandem mass spectrometry

As previously described, peptides were resuspended in 10 μL of 3% acetonitrile, 0.1% formic acid and 1 μL was used for evaluation by liquid chromatography (LC) coupled tandem MS (LC-MS/MS) using an UltiMate^®^ 3000 Rapid Separation LC system (Dionex Corporation, Subbyvale, CA) coupled to a Q Exactive HF mass spectrometer (Thermo Fisher).

### Mass spectrometry data analysis and statistics

Raw data were processed using MaxQuant software (v1.6.17.0) [17] against the human proteome obtained from Uniprot (May 2021) [18]. Variable modifications were set as methionine oxidation and N-terminal acetylation, with a fixed modification of carbamidomethylation of cysteine. The protein and PSM false discovery rates (FDR) were set at 0.01 and “Match between runs” was enabled.

Comparative statistical analysis was performed using MSqRob (v0.7.7) [19] in the R environment (v4.1.0) [20]. Proteins significantly changing between conditional comparisons were taken at 5% FDR. Reactome pathway analysis of differentially expressed proteins was performed using the R package ReactomePA (1.36.0) [21].

### Histological Imaging

Stained slides were imaged using a DMC2900 Leica instrument with Leica Application Suite X software (Leica).

### Data Availability

The mass spectrometry proteomics data have been deposited to the ProteomeXchange consortium via the PRIDE [22] partner repository with the dataset identifier PXD029341.

## Results

### Spatial resolution of the IPF fibrotic front identifies distinct protein signatures

**Figure 1** shows a schematic of our laser capture microdissection approach demonstrating precise cut and capture of the FF, adjacent mature scar, and adjacent alveoli in an IPF specimen. Pentachrome stain identifies immature collagen (highlighting the FF) via the blue color while mature scar tissue appears yellow. We performed LCM on 6 IPF specimens, capturing 3 regions per specimen: IPF alveoli, IPF FF and IPF mature scar (a total of 18 samples). We captured alveoli from 6 non-fibrotic controls (morphologically normal tissue adjacent to tumors during lung resection). For this preparation (a total of 24 samples), we collected tissue volumes of roughly 0.1 mm^3^ per sample which were then processed for mass spectrometry (MS). This procedure involved detergent-based heat-retrieval, physical disruption, and an in-column trypsin digest system to maximize peptide yield, as we did previously [15].

**Figure 1:**
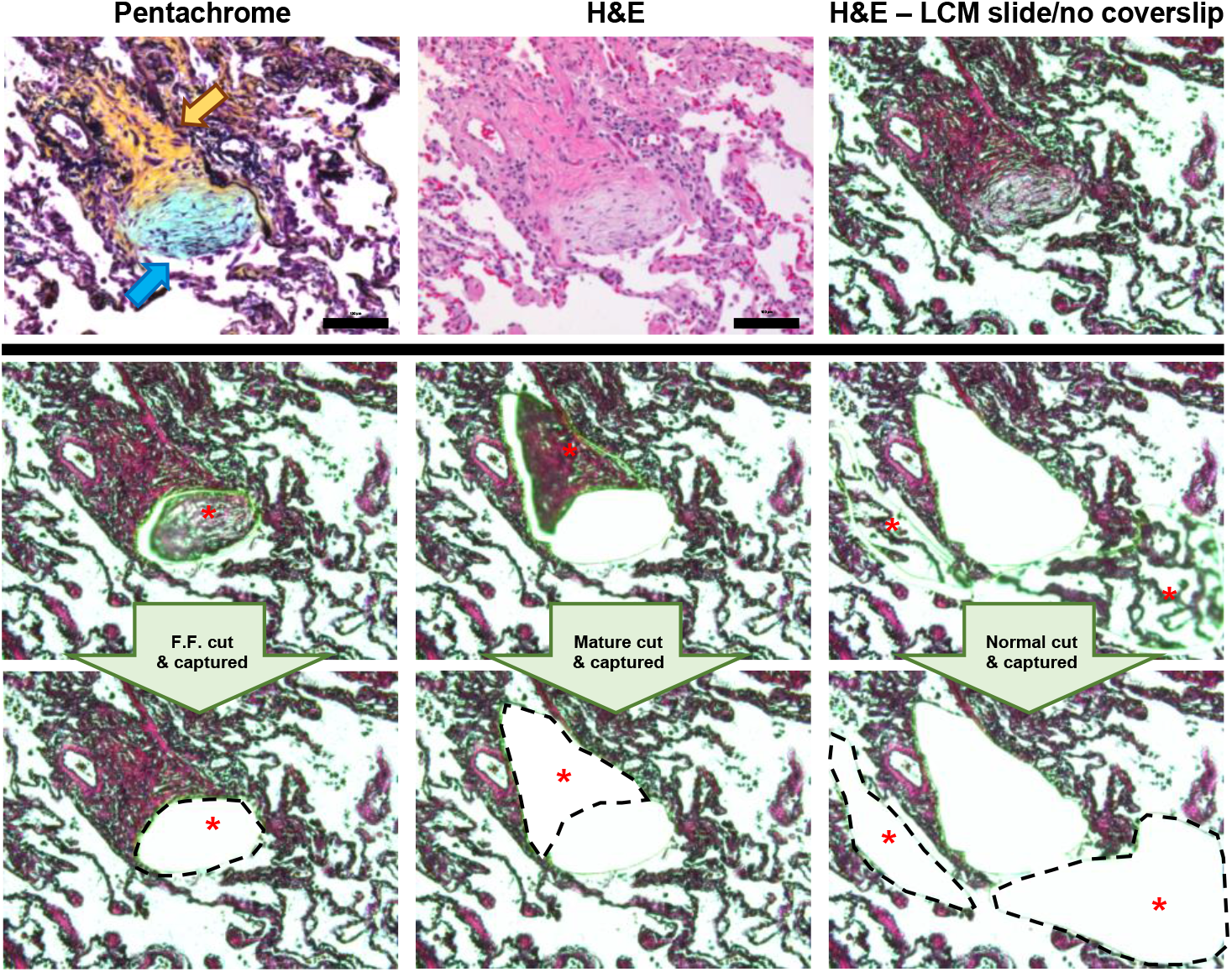
Laser capture microdissection of the fibroblastic focus, mature scar, and adjacent alveoli in an IPF specimen. Formalin-fixed paraffin-embedded specimens were serially sectioned at 5 microns and stained with pentachrome (Top left) or H&E (the other 8 panels). Notice that pentachrome stains the fibroblastic focus (signature lesion of IPF) in the color blue [blue arrow], while the mature scar tissue appears yellow in color [yellow arrow]. We individually captured the fibroblastic focus (left middle and lower panels), the mature scar tissue (mid-middle and lower panels), and adjacent alveoli (right - middle and lower panels) for mass spectrometry preparation and analysis. Scale bar represents 100 microns.

In our qualitative analysis approach, we confirmed the presence of a protein within a sample if it was detected in 3 or more of the 6 samples per group. Using this threshold, qualitative analysis of the MS data showed that we detected 3,147 proteins with the greatest number in non-fibrotic alveoli control (**Figure 2A;** a full list of proteins in **Supplemental File 1**). A 3-dimensional principal component analysis (PCA) based on quantitative MS data (a differential analysis of the proteins using MSqRob v0.7.7 [19]) showed that each region uniquely clusters (**Figure 2B**). Firstly, we saw clear separation of protein signatures in the non-fibrotic alveoli control (top right red dots) versus all of the IPF samples (all other dots on the left), including the structurally intact IPF alveoli (yellow dots). Secondly, we found that the IPF FF cluster (dark green dots) was the most distant from the non-fibrotic alveoli control cluster (red dots). The IPF mature scar is intermediate between the IPF FF and IPF alveoli. The protein signatures thus suggests that the IPF fibrotic front is a distinct environment showing regional changes associated with fibrosis progression.

**Figure 2.**
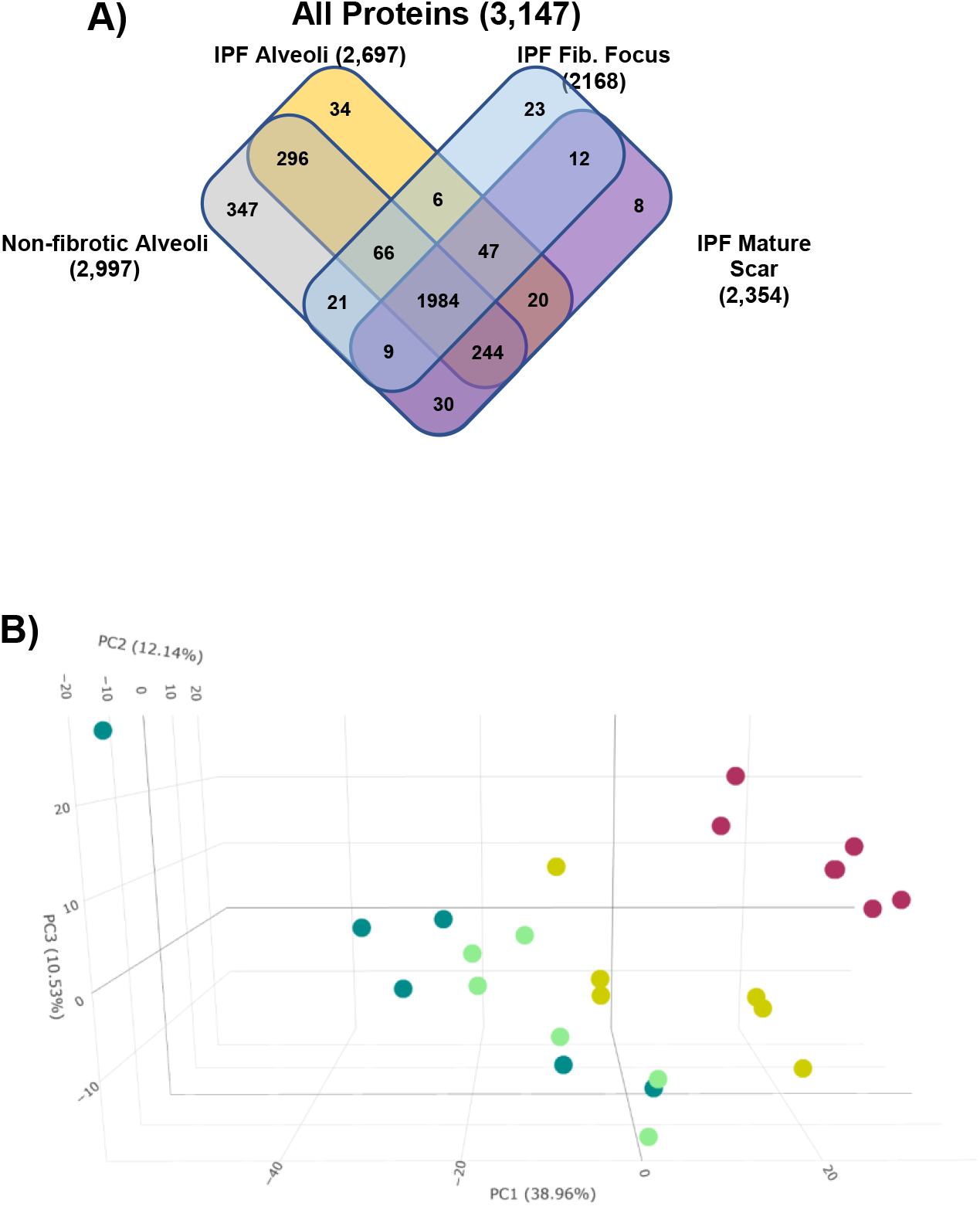
Spatial proteomic analysis of IPF. IPF specimens were subjected to laser capture microdissection coupled mass spectrometry (LCM-MS) to collect Mature Scar, Fibroblastic Focus, and IPF alveoli (n = 6 IPF specimens). In addition, LCM-MS was performed to collect alveoli from non-fibrotic controls (n = 6 non-fibrotic specimens). (**A**) Venn diagram showing all proteins found in each region. (**B**) 3-D principle component analysis showing that the fibroblastic focus (dark green dots) is most distant from non-IPF alveoli (red dots), with the mature scar (light green dots) returning towards IPF Alveoli (yellow dots).

### The alveoli of IPF are enriched with immune-regulatory proteins

Previous reports suggest that the alveoli in IPF are not normal [23–26]; instead, the transition from normal lung to fibrosis shows early pathological features, such as airway remodelling with immune cell infiltration (T-cell, B-cell, and macrophages). We therefore compared IPF alveoli to non-fibrotic alveoli control in more detail. We find that there are 45 proteins that are higher in IPF alveoli versus 211 proteins higher in non-fibrotic alveoli control (**Figure 3A;** a full list in **Supplemental File 2**). Among the proteins downregulated in IPF alveoli was AGER (the gene for the inflammatory receptor RAGE), which was previously shown to be suppressed in IPF lung and is a central mediator of inflammation (high in non-fibrotic alveoli control) [23, 27]. The mitochondrial fusion protein MFN2, which was higher in IPF alveoli, is of interest as its knockout in alveolar type II cells (AECII) was found to induce lung fibrosis [28]. BRK1 (also known as HSPC300), a regulator of the WAVE complex that controls actin assembly [29], is increased in IPF alveoli. The cytoskeletal protein DSP that links intermediate filaments to membrane proteins is also high in IPF alveoli. DSP also shows high expression in IPF airway cells, suggesting a possible role in IPF pathogenesis [30].

**Figure 3.**
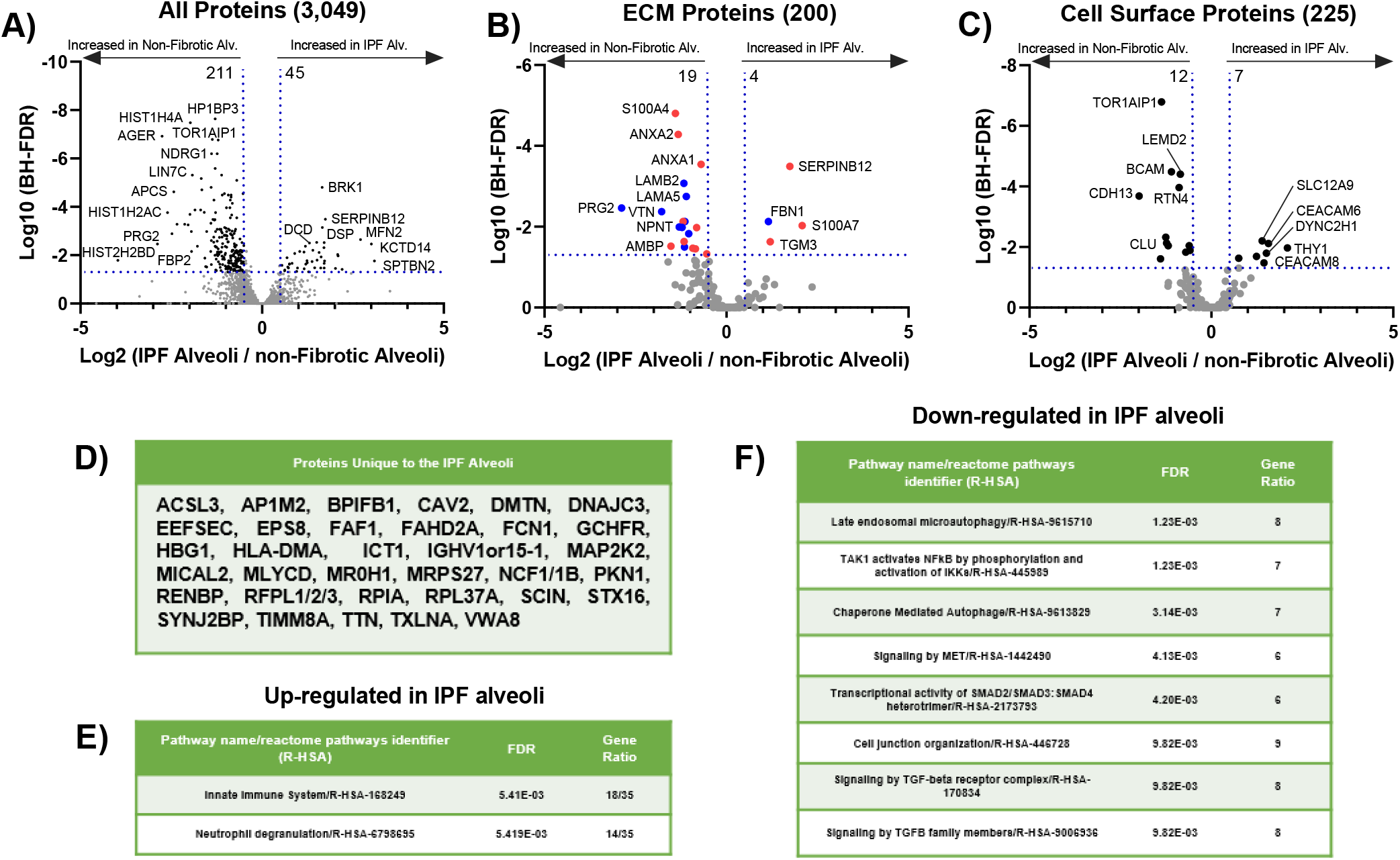
Immune dysregulation defines IPF Alveoli. (**A-C**) Volcano plots comparing IPF alveoli to non-fibrotic alveoli showing the negative natural log of the false discovery values (FDR) values plotted against the base 2 log (fold change) for each protein. The data in (**A**) is for all proteins, whereas (**B**) was matched against the ‘Human Matrisome Project’ (http://matrisomeproject.mit.edu) and (**C**) was matched against the ‘Cell Surface Protein Atlas’ (http://wlab.ethz.ch/cspa/). (**D**) Proteins unique to IPF Alveoli. Reactome Pathways showing the most (**E**) up-regulated or (**F**) down-regulated for IPF alveoli compared to non-fibrotic alveoli.

We further separated our data to identify ECM proteins (**Figure 3B**) that differ between IPF alveoli and non-fibrotic alveoli control by matching our data set to the ‘Human Matrisome Project’ (http://matrisomeproject.mit.edu) [31]. IPF Alveoli have higher levels of transglutaminase 3 (TGM3). This protein crosslinks components of the extracellular matrix, which increases matrix rigidity but has also been linked to EMT and Akt signalling pathway in colorectal cancer [32]. The transmembrane protein PRG2 was implicated in neuronal growth by interacting with and modulating phosphatase and tensin homolog deleted on chromosome 10 (PTEN) [33], and is lower in IPF alveoli. PTEN is a potent negative regulator of PI3-lipids and Akt, which promote both cell invasion and survival. Akt signalting is activated in IPF [34, 35], thus, the concept that changes in the ECM might drive this pathway warrants further study.

Utilizing the ‘Cell Surface Protein Atlas’ (http://wlab.ethz.ch/cspa/) [36], we matched our dataset to the ‘Surfaceome’ to determine which cell surface receptors are altered in different regions of the IPF lung (**Figure 3C**). Both the cell-cell adhesion protein CEACAM6 and the GPI-linked protein Thy1 are increased in IPF alveoli. Interestingly, both are involved in immune cell regulation and are over-expressed in malignancies [37, 38]. Thy1 is also implicated in regulating integrin function in the context of fibrosis [39]. Interestingly, TOR1AIP1, a protein of the nuclear membrane, is also decreased in IPF alveoli. Variants of this gene have been identified in limb-girdle muscular dystrophy and TOR1AIP1 knockout in striated muscle results in muscle weakness [40, 41]. Our qualitative data identify 34 proteins detected only in IPF alveoli (**Figure 3D**) whereas 347 proteins are unique to non-fibrotic alveoli control (**Supplemental File 3**). ACSL3, which is unique to the IPF alveoli, is elevated in pancreatic ductal carcinoma where it correlates with increased fibrosis [42].

We next used the unbiased Reactome pathway analysis to determine which pathways are perturbed in IPF alveoli. The two significantly upregulated in IPF alveoli were ‘Innate Immune System’ and ‘Neutrophil degranulation’ (**Figure 3E**), consistent with immune involvement in progression of fibrosis [43]. Down-regulated pathways include ‘Late endosomal microautophagy’, several categories related to TGFβ signaling, and ‘Cell junction organization’ (**Figure 3F**). Downregulated TGFβ signaling at this early, pre-fibrotic stage would be consistent with the elevated inflammatory status. These data suggest that early events towards IPF progression occur well before morphological changes are evident.

### The IPF mature scar is consistent with end-stage fibrosis

Comparison of the IPF mature scar to non-fibrotic alveoli control identified 81 proteins increased and 651 proteins decreased in IPF mature scar relative to non-fibrotic alveoli control (**Figure 4A;** a full list in **Supplemental File 2**). The endoplasmic reticulum chaperone protein MZB1 was highly elevated in IPF mature scar, consistent with a report that MZB1-positive plasma B cells (that co-express CD38) are found in end-stage lung and skin fibrosis [44]. Supporting possible B cell involvement, CD38 was among the uniquely expressed proteins in IPF mature scar tissue **(Figure 4D)**. The epithelial polarity and scaffolding protein LLGL2 was decreased in IPF mature scar, consistent with its high expression in polarized epithelial cells and its loss during EMT [45].

**Figure 4.**
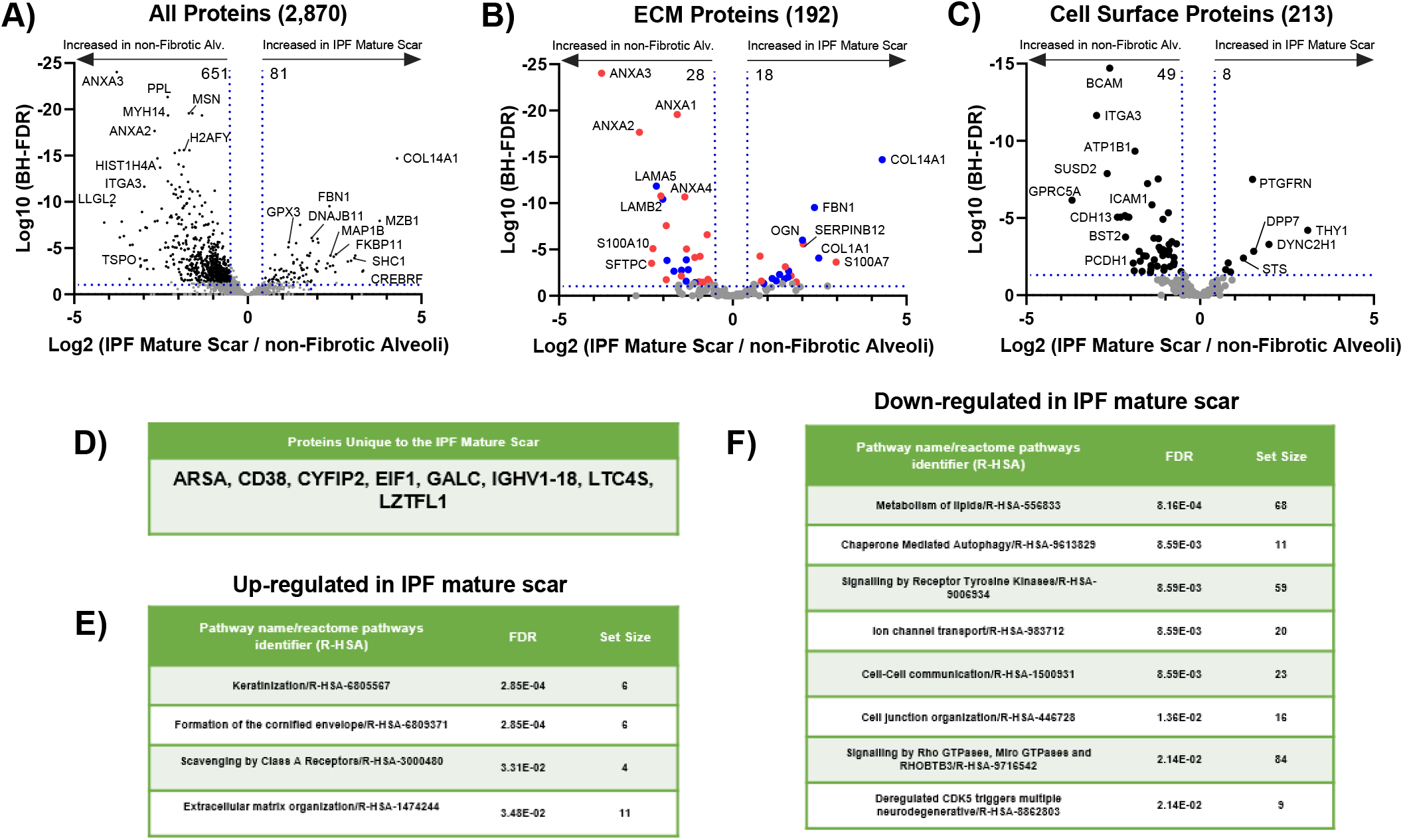
End-stage fibrosis defines IPF Mature Scar. (**A-C**) Volcano plots comparing IPF mature scar to non-fibrotic alveoli showing the negative natural log of the false discovery values (FDR) values plotted against the base 2 log (fold change) for each protein. The data in (**A**) is for all proteins, whereas (**B**) was matched against the ‘Human Matrisome Project’ (http://matrisomeproject.mit.edu) and (**C**) was matched against the ‘Cell Surface Protein Atlas’ (http://wlab.ethz.ch/cspa/). (**D**) Proteins unique to IPF mature scar. Reactome Pathways showing the most (**E**) up-regulated or (**F**) down-regulated for IPF mature scar compared to non-fibrotic alveoli.

We again matched our dataset with the ‘Matrisome project’ (**Figure 4B**). Multiple annexin family proteins were decreased in IPF mature scar. ANXA1 has anti-inflammatory effects and its loss-of-function exacerbates inflammation in bleomycin-induced lung injury models [46]. ANXA3 is decreased in IPF but reportedly promotes inflammation [47], whereas ANXA2 is anti-inflammatory [48]. Among collagen proteins, Col14a1 was the most highly upregulated in IPF mature scar. This gene was also a marker for a fibroblast cluster found in bleomycin-induced lung injury by single cell sequencing [49]. Interestingly, collagen XIV is elevated in high mechanical stress environments where it regulates fibrillogenesis [50].

We also matched our dataset to the ‘Surfaceome’ (**Figure 4C**). The transmembrane Ig superfamily protein PTGFRN was strongly increased in IPF mature scar. This protein was previously shown to be increased in both IPF and in bleomycin-induced lung injury [51]. By contrast, the laminin receptor BCAM, was decreased in IPF mature scar, consistent with decreased expression of laminin proteins in this region [52]. Reactome pathway analysis demonstrates that the categories most strongly upregulated in IPF mature scar were ‘Keratinization’ and ‘Formation of the cornified envelope’ (**Figure 4E**), whereas ‘Metabolism of lipids’ was most strongly decreased (**Figure 4F**). These changes likely reflect the heavily cross-linked, rigid structure and low metabolic activity of end-stage fibrosis [53].

### The IPF fibroblastic focus shows elevated collagen biosynthesis

A previous study using laser capture microdissection of the fibroblastic focus coupled with RNA-sequencing and a novel weighted gene co-expression network analysis showed elevation of genes associated with cell cycle, inflammation/differentiation, translation, and cytoskeleton/cell adhesion [54]. Our analysis identified 100 proteins increased and 211 decreased in the IPF FF relative to non-fibrotic alveoli control (**Figure 5A;** a full list in **Supplemental File 2**). These data were also matched to both the ‘Matrisome project’ and the ‘Surfaceome’ (**Figure 5B – 5C**); the 22 proteins uniquely expressed in the IPF fibroblastic focus are shown in **Figure 5D**.

**Figure 5.**
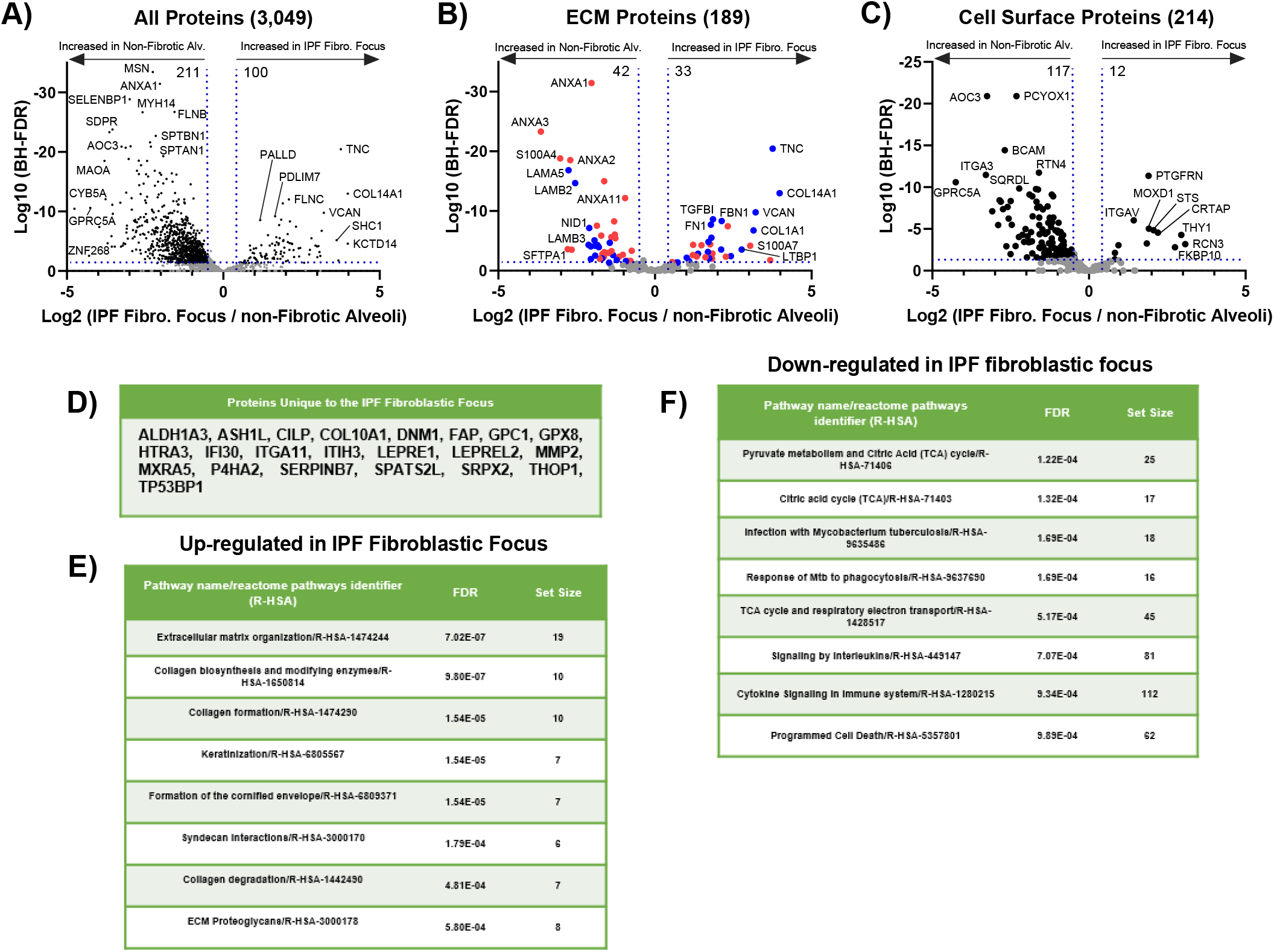
The fibroblastic focus is an active collagen biosynthesis factory. (**A-C**) Volcano plots comparing the IPF fibroblastic focus to non-fibrotic alveoli showing the negative natural log of the false discovery values (FDR) values plotted against the base 2 log (fold change) for each protein. The data in (**A**) is for all proteins, whereas (**B**) was matched against the ‘Human Matrisome Project’ (http://matrisomeproject.mit.edu) and (**C**) was matched against the ‘Cell Surface Protein Atlas’ (http://wlab.ethz.ch/cspa/). (**D**) Proteins unique to the IPF fibroblastic focus. Reactome Pathways showing the most (**E**) up-regulated or (**F**) down-regulated for the IPF fibroblastic focus compared to non-fibrotic alveoli.

Collagen biosynthesis is a complex, multistep process that involves collagen hydroxylation, triple helix formation, crosslinking/maturation, secretion via a specific pathway and uptake/turnover (reviewed in [55]). Collagen is also post-translational modified by prolyl-4-hydroxylases (P4HA proteins) and lysyl hydroxylases (PLOD proteins) to promote its stability and function. Consistent with collagen post-translational modifications (PTMs), the prolyl-4-hydroxylase P4HA2 is unique to the IPF fibroblastic focus. The lysyl hydroxylase PLOD2 is unique to both the IPF fibroblastic focus and IPF mature scar, while PLOD1 is significantly increased in the IPF fibroblastic focus.

The next step in collagen biosynthesis is the formation of the triple helix. FKBP10, which plays a critical role in triple helix formation, is increased in the FF, as is SerpinH1 (Hsp47), which functions as an essential collagen-specific chaperone needed for collagen synthesis via effects on triple helix formation and modification [55, 56]. In addition, the prolyl 4-hydroxylase LEPREL2, a gene required for proper collagen folding and assembly [57], is unique to the IPF FF. Collagen is crosslinked/matured by a host of enzymes such as the transglutaminase (TGM) family. TGM1 and TGM3 are increased in the IPF FF. Lastly, collagen uptake is mediated by the integrin subunits alpha-1, −2, −10, −11 and beta-1, whereas collagen cleavage is mediated by matrix metalloproteinases (MMPs). MMP14 was increased and MMP2 uniquely expressed in the IPF FF. Integrin alpha 11 subunit (ITGA11), a collagen I specific receptor, is unique to the IPF FF, while integrin alpha-1 and beta-1 are decreased. The consequences of these changes are unknown but effects on collagen fibril organization or turnover are likely.

Reactome pathway analysis shows that the IPF FF is indeed enriched for ‘Collagen biosynthesis and modifying enzymes’, ‘Collagen formation’ and ‘ Collagen degradation’ (**Figure 5E**) which is in accord with prior data showing that the fibroblastic focus uniquely stains with pro-collagen (active collagen synthesis) [16, 58, 59]. Interestingly, ‘pyruvate metabolism and Citric Acid (TCA) cycle’ were down-regulated in the IPF FF, which may correlate with low cell proliferation in this region (**Figure 5F**). These data strongly suggest that the IPF fibroblastic focus is the major site of active collagen biosynthesis.

### The IPF fibroblastic focus experiences a dramatic extracellular matrix switch

We next aligned all of our data (IPF FF, IPF mature scar, IPF alveoli, and non-fibrotic alveoli control) to derive a heatmap of all statistically significant ECM proteins matched to the 6 matrisome categories (Glycoproteins, Proteoglycans, ECM-affiliated proteins, ECM Regulators, Secreted Factors, and Collagens) (**Figure 6A**). In every case, the ECM proteins comprising the IPF FF (far left) show the greatest difference from non-fibrotic alveoli control (far right). For instance, it is well known that collagen IV, a major constituent of epithelial basement membranes, decreases in IPF [60]. We also found that collagen IV is low in both the IPF FF and IPF mature scar, and high in both IPF alveoli and non-fibrotic alveoli control. Similarly, collagen I is high in both IPF mature scar and IPF FF, and low in both IPF alveoli and non-fibrotic alveoli control [in lowest panel ‘collagen’ of **Figure 6A**]. Overall, IPF alveoli tend to resemble the non-fibrotic alveoli controls, whereas IPF mature scar tends to resemble the IPF fibroblastic focus, with substantial deviations. Examination of the qualitative data to determine the numbers of ECM proteins by categories and per region [**Supplemental Figure 1**] also revealed no dramatic difference in the types of ECM proteins found within each region. Taken together, these results reveal a dramatic switch of ECM in the IPF fibroblastic focus.

**Figure 6.**
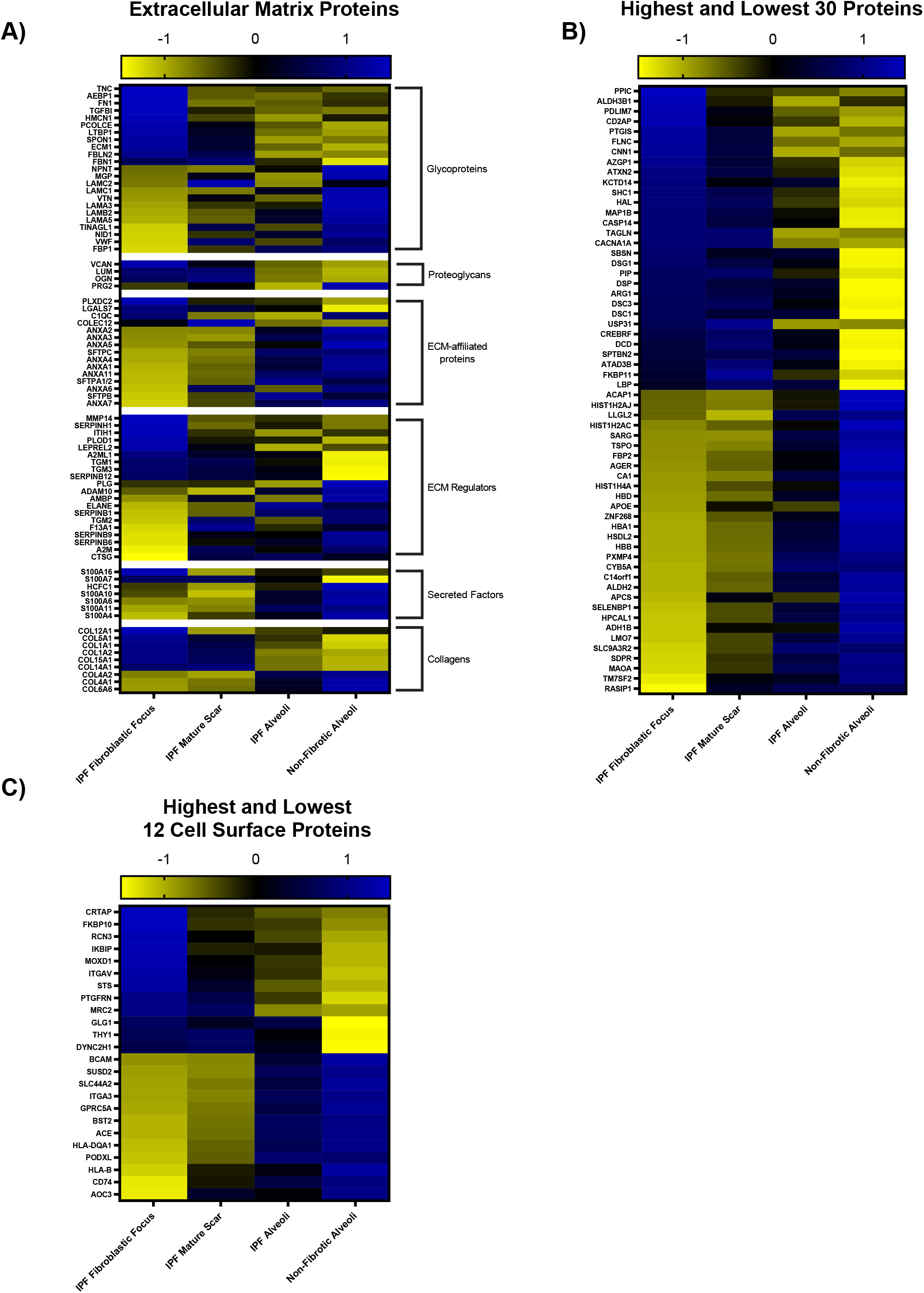
The IPF fibroblastic focus has a unique ECM signature. Shown are heatmaps to demonstrate the expression of (**A**) ECM proteins, (**B**) the highest and lowest 30 proteins (excluding ECM and cell surface proteins), and (**C**) highest and lowest 12 cell surface proteins in the IPF fibroblastic focus, IPF mature scar, IPF alveoli, and non-fibrotic alveoli. Blue indicates protein is increased and yellow indicates the protein is decreased.

Similarly, we created a heatmap of the highest and lowest 30 statistically most changed proteins (**Figure 6B;** a full list in **Supplemental File 4**). The top 3 proteins expressed in the IPF fibroblastic focus are PPIC, ALDH3B1, and PDLIM7. PPIC is associated with protein folding and is upregulated in CCL4-induced liver injury. PPIC loss-of-function was shown to improve liver injury suggesting it may be involved in fibrosis progression [61]. Similarly, ALDH3B1, which is involved in oxidative stress, is highly expressed in lung cancer [62]. PDLIM7 regulates cellular senescence [63]; a process that likely contributes to IPF [64]. Analysis of the highest and lowest 12 statistically changed cell surface proteins (**Figure 6C**) identified CRTAP and RCN3 as the most increased in the IPF FF. Both genes are critical for collagen biosynthesis [65, 66]. In accord with reports showing that the IPF fibroblastic focus is hypoxic [67], the IPF FF showed increased steroid sulfatase (STS), a gene involved in steroid hormone synthesis that was reported to induce hypoxia inducible factor 1 (HIF1) expression [68]. We also noted that both integrin alpha-v (ITGAV; cell surface protein) and latent transforming growth factor beta-binding protein 1 (LTBP1; ECM protein) increased in the IPF FF, consistent with their roles in activating latent TGF-β [69]. This study provides us with a list of some established and many new region-specific proteins to begin exploring their collective functions in the pathogenesis of IPF.

Lastly, we sought to compare the IPF fibroblastic focus with IPF mature scar tissue (**Supplemental Figure 2**, a full list in **Supplemental File 5**). The fibroblastic focus shows greater up-regulation of proteins involved in collagen biosynthesis and ECM organization. Major ECM proteins highlighted here include TNC, CTHRC1, SERPINH1, FN1, VCAN, and COL12A1, all of which are involved in assembly or stabilization of the fibrotic ECM.

To validate some of these results, we immunostained for SERPINH1 and COL12A1 in 4 IPF specimens and 2 non-fibrotic controls. Consistent the proteomic analysis, we found both specifically enriched within the FF (**Figure 7A;** additional images in **Supplementary Figure 3**). Expression of SERPINH1 (essential for collagen synthesis) within the fibroblastic focus was consistent with its role in collagen biosynthesis. By contrast, neither SERPINH1 nor COL12A1 were detected in non-fibrotic controls (**Figure 7B**).

**Figure 7.**
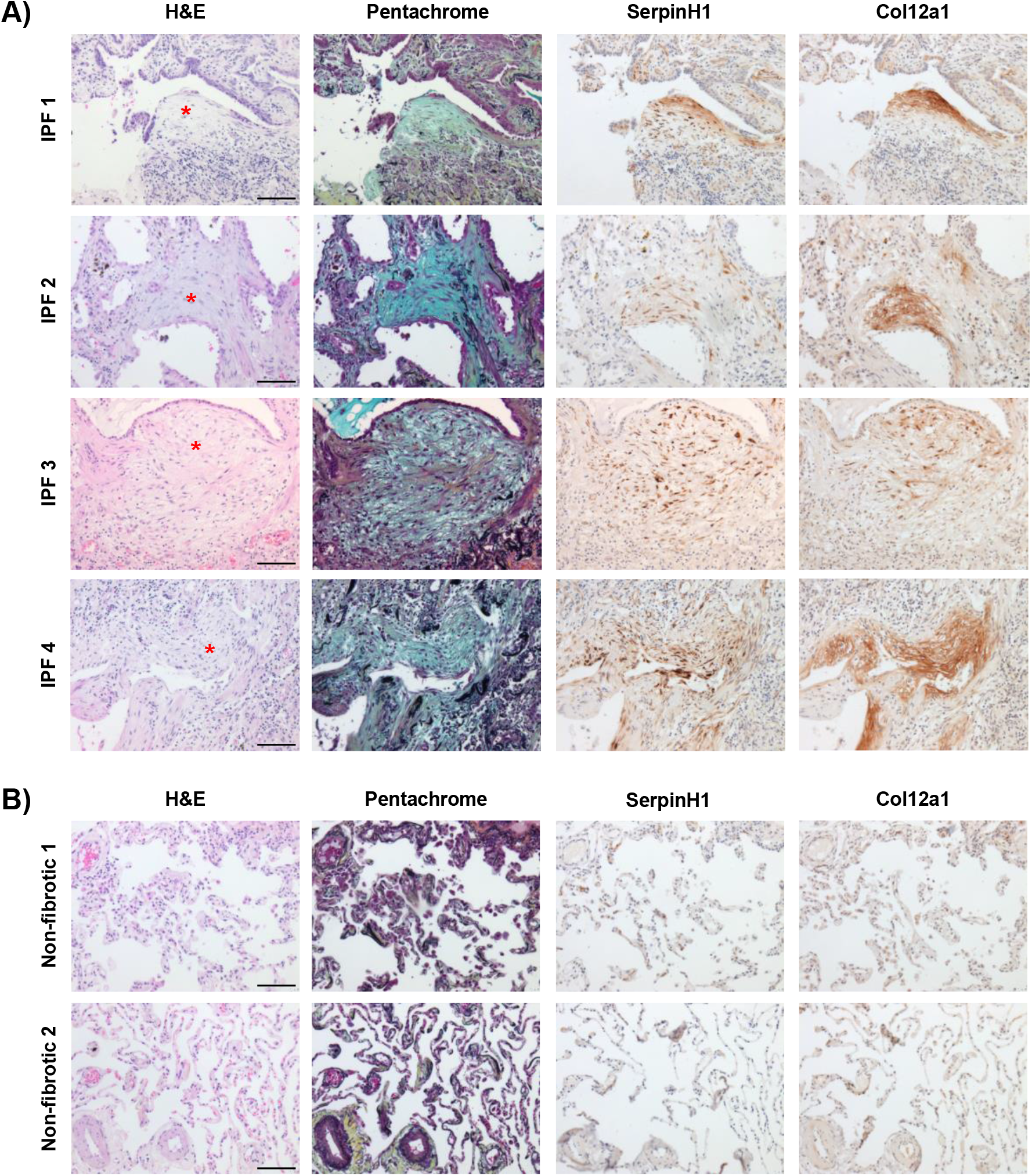
SerpinH1 and Col12a1 are enriched in the fibroblastic focus. 4 IPF specimens were serially sectioned and stained for H&E, pentachrome, anti-SerpinH1, and anti-Col12a1. We show on representative fibroblastic focus per specimen (depicted with a red asterisk). Scale bar represents 100 microns.

## Discussion

The availability of spatial ‘omics’ makes it possible to reveal previously inaccessible clinical features of human disease. Herein, we produced spatial proteomic profiles of the IPF fibrotic front to characterize the protein composition of non-fibrotic alveoli, IPF alveoli, the IPF fibroblastic focus, and IPF mature scar. We created a tissue atlas of the IPF fibrotic front (**Figure 8**) defining the ECM and cell surface proteins of each region in an effort to inform novel hypotheses about cell/ECM systems in IPF progression. Recent work utilizing atomic force microscopy to define mechanical properties of the different zones of IPF tissue showed that the IPF fibroblastic focus (2.0 kPa) is surprisingly as soft as adjacent IPF alveoli (1.5 kPa), whereas the IPF mature scar is much stiffer (9.0 kPa) [70]. Unfortunately, non-fibrotic controls were not included in that report. The low stiffness of the fibroblastic focus was surprising, especially in light of our finding of increased levels of a variety of collagen and ECM proteins. We speculate that the enrichment of glycoproteins and proteoglycans, which have the capacity to hydrate tissues [71], may be responsible for the softness of the FF.

**Figure 8.**
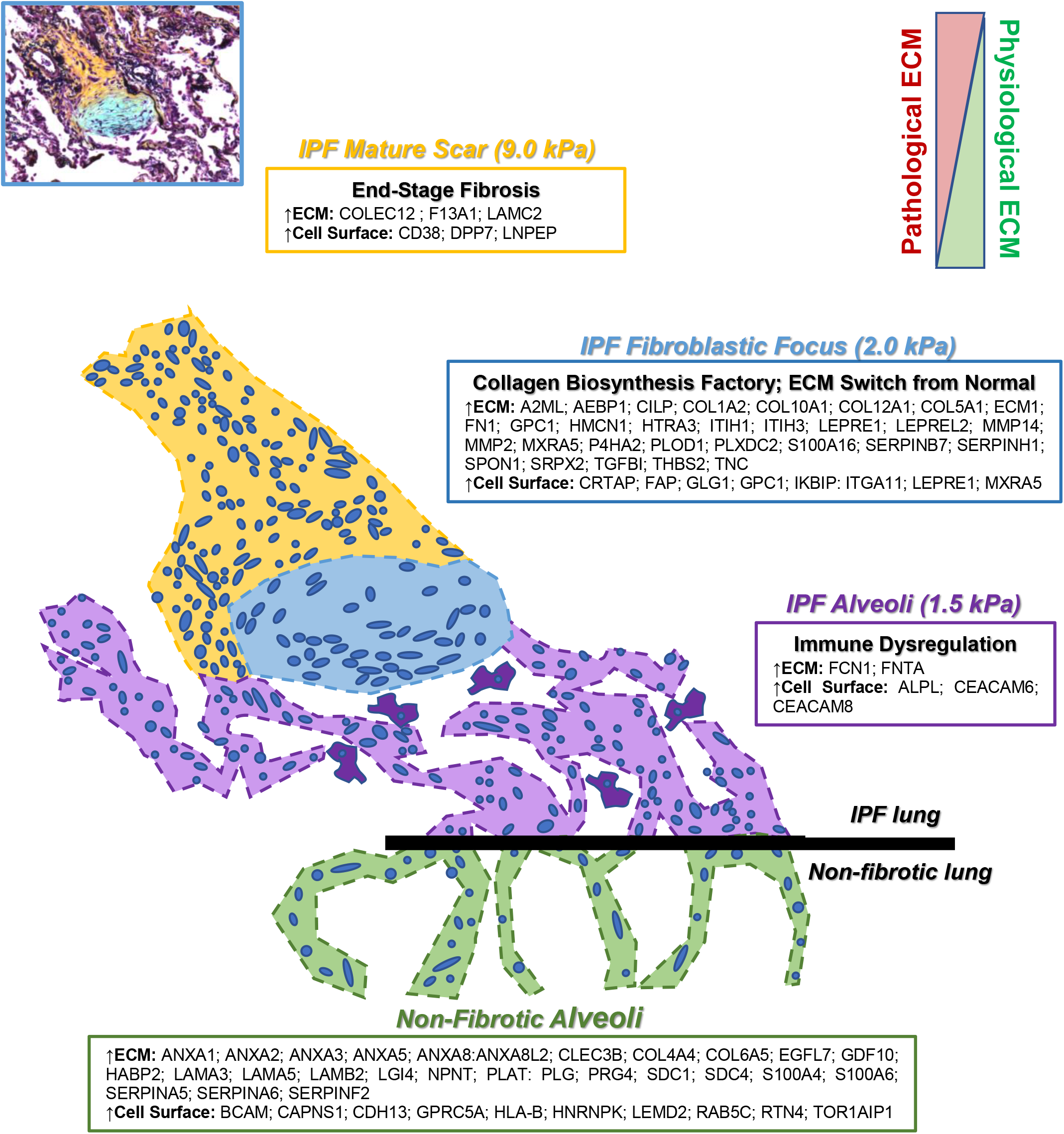
The Idiopathic Pulmonary Fibrosis fibrotic front. The lesion of IPF is termed the fibroblastic focus (FF) and can be envisioned as the invasive front. The FF progresses towards adjacent alveoli leaving behind a dense mature scar tissue. Herein, we provide a list of proteins (extracellular matrix and cell surface) that are abundantly and significantly over-expressed per region. We define the fibroblastic focus as a collagen biosynthesis factory that is embedded in a unique extracellular matrix; a switch from normal. Adjacent to the fibroblastic focus, the mature scar, is characterized as end-stage fibrosis which is stiffer than the rest of the IPF tissue. The alveoli adjacent to the fibroblastic focus is defined by immune dysregulation.

Convincing evidence suggests that normal-appearing alveoli within the IPF lung are in fact significantly abnormal. RNA sequencing of tissue from different regions of the IPF lung, showed that structurally normal IPF tissue had over 1,000 differentially expressed genes compared to non-fibrotic tissue [23]. Increased immune cell infiltration (innate and adaptive) has also been observed in structurally normal regions of IPF lung [24]. 3-dimensional image analysis of the IPF lung (specifically looking at regions with no evidence of microscopic fibrosis) showed reduced numbers of small airways and thickening of terminal bronchiolar walls, suggesting that early remodelling is crucial to IPF pathogenesis [26]. One important unanswered question is whether the changes of the alveoli observed here indicated early steps in IPF development, which eventually lead to the development of the fibroblastic focus and then mature scar tissue. Alternatively, it may be that the fibroblastic focus and mature scar affect surrounding lung tissue, potentially through secreted factors that act directly, by promoting immune cell infiltration, or by altering lung mechanics. Future studies analysing spatial proteomics in multiple regions of the uninvolved IPF lung are needed to address this question.

The mature scar region in IPF is consistent with end-stage fibrosis. McDonough et al. similarly sampled regions of the IPF lung from structurally normal to increasing fibrosis (IPF level 1 through 3). They concluded that at the transcriptional level, the transition from level 2 to level 3 (the most fibrotic tissue) involves genes characteristic of end-stage disease [23], consistent with the higher stiffness in these regions [70]. This feature is diagnostic of both denser and more crosslinked collagen, as suggested by higher levels of transglutaminase and lysyl hydroxylase. These results suggest that critical pathways within the fibroblastic focus are the most promising targets for inhibiting progression of IPF.

The IPF fibroblastic focus is the signature lesion of IPF [1] and is consistently described as the site of collagen synthesis. Our data defining the protein constituents of this domain lead to two major conclusions. First, a large fraction of the proteins necessary for collagen biosynthesis are either increased or uniquely expressed within the fibroblastic focus. The fibroblastic focus is therefore the primary region of active collagen biosynthesis. Second, we provide evidence that the ECM of the fibroblastic focus is completely switched from non-fibrotic alveoli controls, with near complete replacement of the epithelial basement membrane with an interstitial collagenous ECM. We speculate that the synthesis and assembly of matrix within the FF may be critical for migration of myofibroblasts or their precursor into uninvolved airspaces. Local inhibition of collagen biosynthesis machinery may therefore be a pharmaceutical approach to stop IPF progression.

## Conclusion

Herein we create an unbiased tissue atlas of the IPF fibrotic front utilizing laser capture microdissection coupled mass spectrometry. We demonstrate that there are regional changes in protein signature associated with IPF progression and provide detailed lists of proteins that suggest hypotheses concerning IPF pathology and progression. The finding that uninvolved alveoli within the IPF lung are highly abnormal suggests that this region may contain novel targets for therapeutic intervention.

## Author Contributions

J.A.H. and M.A.S. conceived and supervised the project. J.A.H. designed and conducted all LCM-MS experiments and C.L. performed the associated analyses. L.D. performed the immunohistochemistry. M.A.M. assisted in characterizing the histological stains and identification of clinical morphologies associated with Idiopathic Pulmonary Fibrosis. R.S., R.V.V. and J.F.B. contributed to reagents. J.A.H. and M.A.S. wrote the manuscript with all author inputs.

## Acknowledgements

The authors would like to thank both Professors Peter Bitterman and Craig Henke for their support in reading and editing the manuscript. We also thank the Histology, BioMS, and BioImaging Facilities at University of Manchester for making this work possible. This report is independent research supported by the North West Lung Centre Charity and National Institute for Health Research Clinical Research Facility at Manchester University NHS Foundation Trust. The views expressed in this publication are those of the author(s) and not necessarily those of the NHS, the North West Lung Centre Charity, National Institute for Health Research or the Department of Health. The authors would like to acknowledge the Manchester Allergy, Respiratory and Thoracic Surgery Biobank and the North West Lung Centre Charity for supporting this project. In addition we would to thank the study participants for their contribution.

## Conflict of Interests

The authors have declared that no conflict of interests exists.

## Funding

This was work was supported by the Wellcome Centre for Cell-Matrix Research’s directors discretional funds (WCCMR; 203128/Z/16/Z) to JAH, Medical Research Council transition support (MR/T032529/1) to JFB, and NIH grant R01 HL135582 to MAS.

---

**Supplemental Figure 1:**
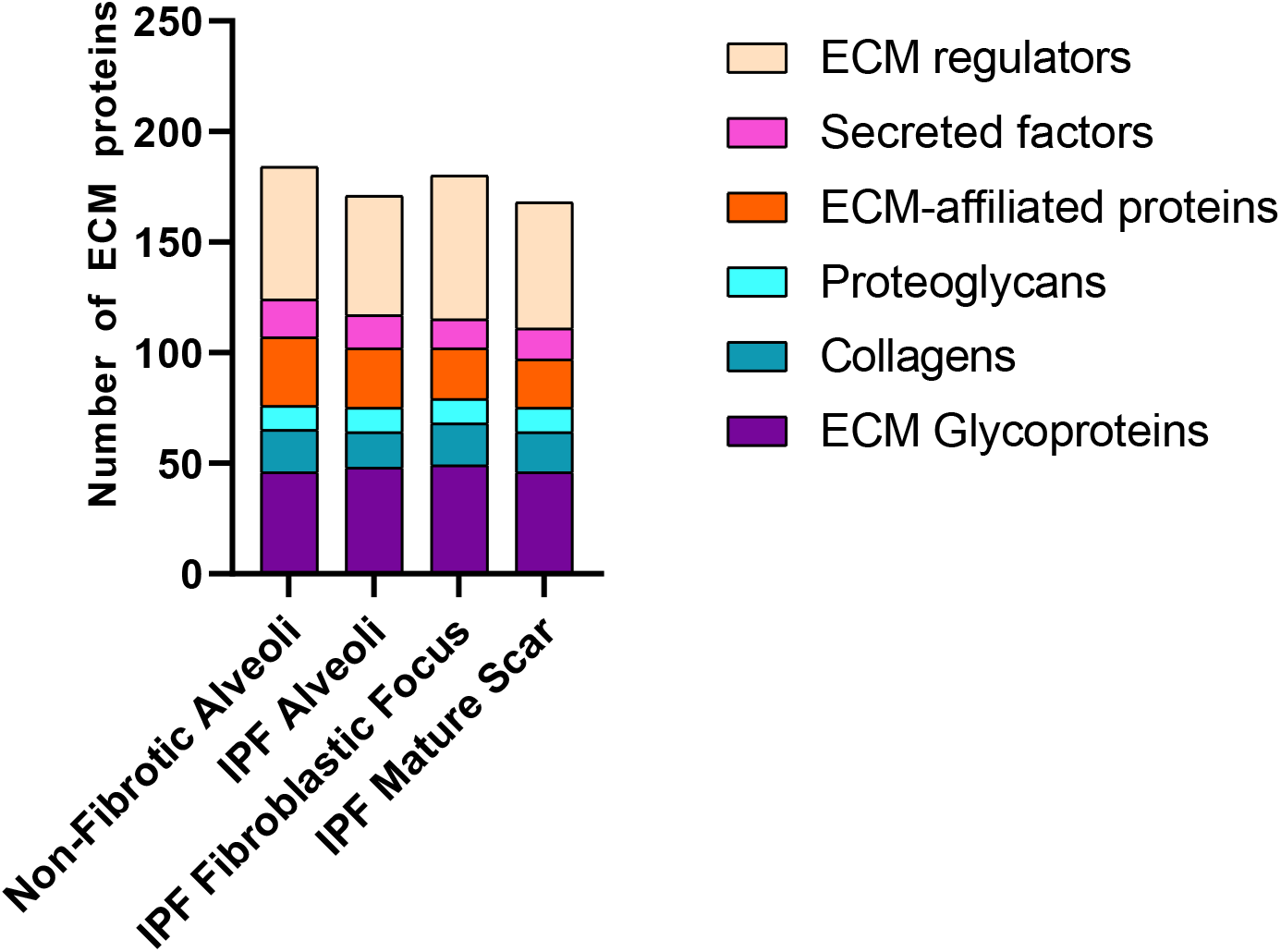
Spatial ECM diversity in IPF. The detected ECM proteins, in each group, were stratified and graphed into 6 ECM categories: ECM glycoproteins, collagens, proteoglycans, ECM-affiliated proteins, ECM regulators, and secreted factors.

**Supplemental Figure 2.**
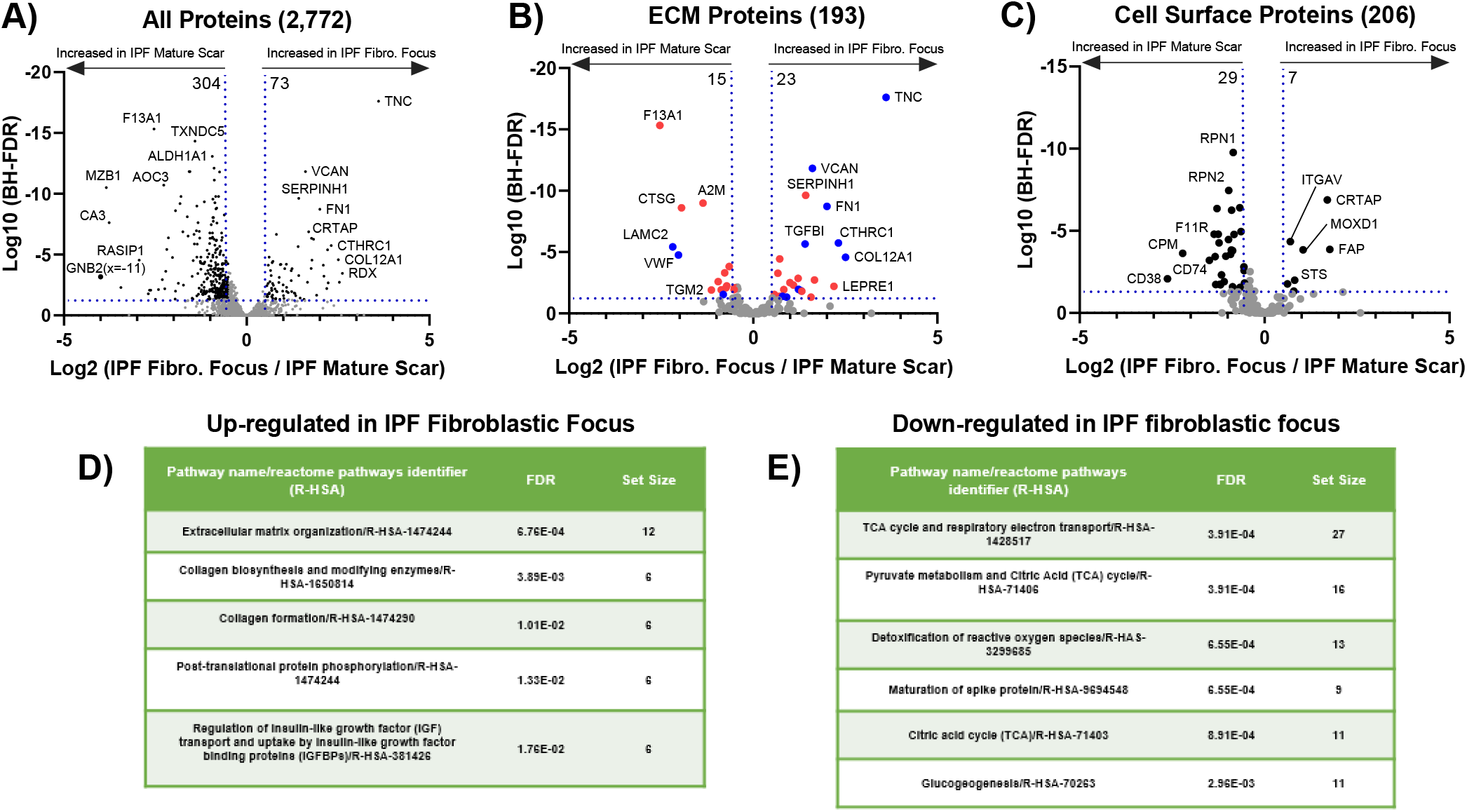
The IPF fibroblastic focus as compared to IPF mature scar tissue. (**A-C**) Volcano plots comparing the IPF fibroblastic focus to IPF mature scar showing the negative natural log of the false discovery values (FDR) values plotted against the base 2 log (fold change) for each protein. The data in (**A**) is for all proteins, whereas (**B**) was matched against the ‘Human Matrisome Project’ (http://matrisomeproject.mit.edu) and (**C**) was matched against the ‘Cell Surface Protein Atlas’ (http://wlab.ethz.ch/cspa/). (**D**) up-regulated or (**E**) down-regulated Reactome pathways for the IPF fibroblastic focus compared to IPF mature scar.

**Supplemental Figure 3.**
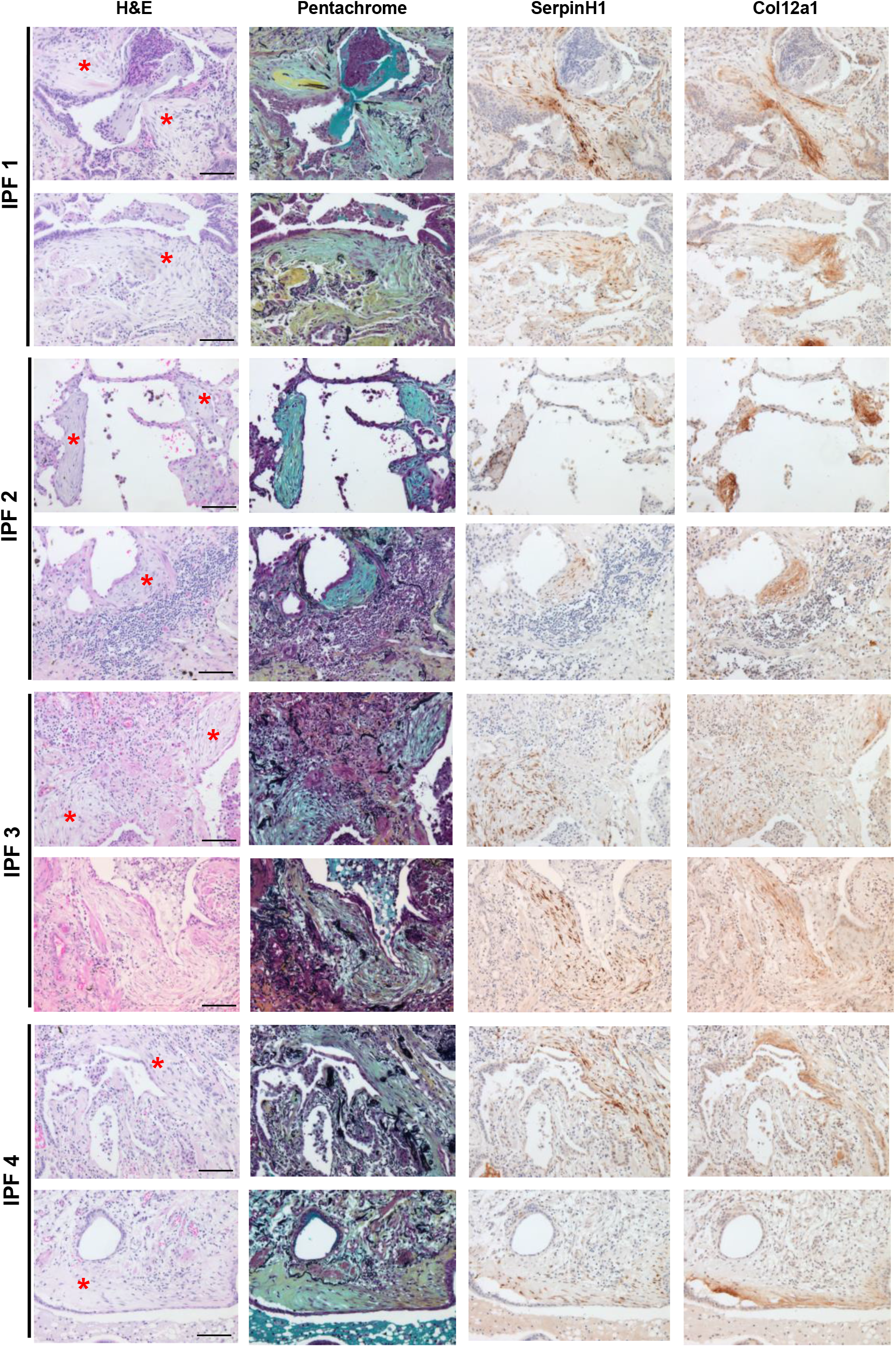
SerpinH1 and Col12a1 expression in IPF. 4 IPF specimens were serially sectioned and stained for H&E, pentachrome, anti-SerpinH1, and anti-Col12a1. We show two additional fibroblastic foci for each specimen (depicted with a red asterisk). Scale bar represents 100 microns.

## Notes

### Competing Interest Statement

The authors have declared no competing interest.

